# PREDICTING GENE DISEASE ASSOCIATIONS WITH KNOWLEDGE GRAPH EMBEDDINGS FOR DISEASES WITH CURTAILED INFORMATION

**DOI:** 10.1101/2024.01.11.575314

**Authors:** Francesco Gualdi, Baldomero Oliva, Janet Piñero

## Abstract

Knowledge graph embeddings (KGE) are a powerful technique used in the biological domain to represent biological knowledge in a low dimensional space. However, a deep understanding of these methods is still missing, and in particular the limitations for diseases with reduced information on gene-disease associations. In this contribution, we built a knowledge graph (KG) by integrating heterogeneous biomedical data and generated KGEs by implementing state-of-the-art methods, and two novel algorithms: DLemb and BioKG2Vec. Extensive testing of the embeddings with unsupervised clustering and supervised methods showed that our novel approaches outperform existing algorithms in both scenarios. Our results indicate that data preprocessing and integration influence the quality of the predictions and that the embeddings efficiently encodes biological information when compared to a null model. Finally, we employed KGE to predict genes associated with Intervertebral disc degeneration (IDD) and showed that functions relevant to the disease are enriched in the genes prioritized from the model

**GRAPHICAL ABSTRACT:** 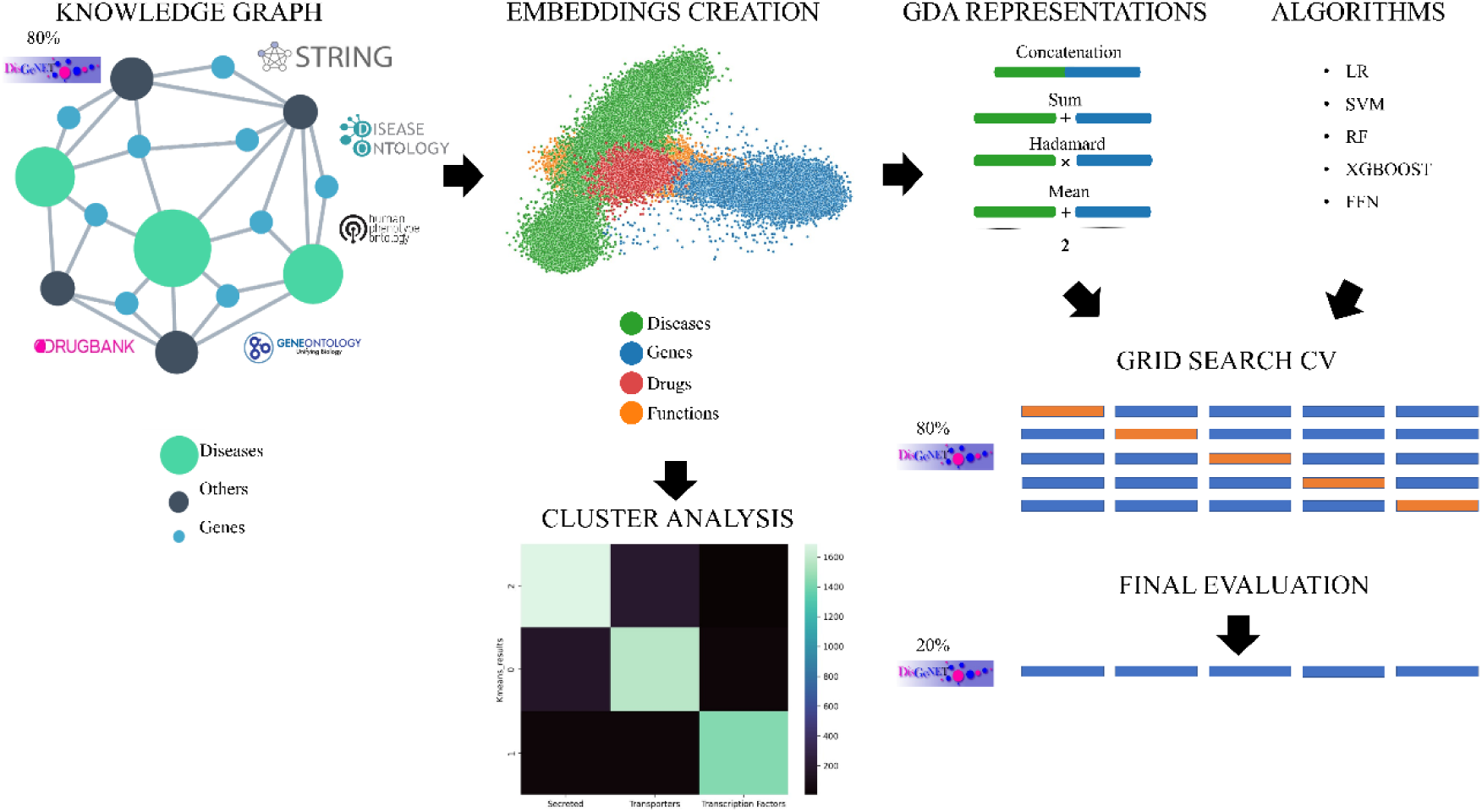

## INTRODUCTION

Predicting genes associated with diseases is a challenging task. Recent advancements in genomic technologies have contributed to reach a deeper understanding of the genetics underlying complex diseases. However, the difficulties related to costs and time of these technologies have prompted the development of *in silico* methods to perform this task (1).

In this regard, network approaches have emerged as valuable tools for building meaningful models allowing the integration of heterogeneous biological knowledge from numerous sources (2). These heterogeneous networks are defined knowledge graphs (KGs), which contain structured depictions of biological systems wherein different biological entities interact through complex relationships. Elucidating these intricate relations is crucial to better interpret complex biological data and thus the possible causes of diseases.

Various methods have been employed to leverage KGs to infer new possible interactions between biological entities. One commonly used approach involves expressing the entities within KGs as low-dimensional vectors using vectorial representations that preserve the graph local structure known as knowledge graph embedding (KGE). This method outperforms other approaches in terms of accuracy and scalability of their prediction (3).

Numerous methods have been developed to generate embeddings from KGs, and they can be broadly categorized into four main families: translational models, semantic matching, random walks-based models, and deep neural networks. Refer to (4, 5) for a comprehensive overview of these methods. Several studies have been conducted to explore the potential of KGE for predicting gene-disease associations (GDAs). For instance, *Nunes et al* investigated the impact of employing rich semantic representations based on more than one ontology to predict GDAs by testing different embedding creation models and machine learning algorithms (6). Other works have focused on the heterogeneous integration of knowledge bases with the development of a single deep learning framework for predicting GDAs starting from a KG (7, 8).

While previous studies have made progress in implementing KGE methods in biomedical research, we lack a proper benchmark of available methods. Existing works in this field are limited to benchmarking the proposed method (7) or the comparison of different algorithms (6) without providing a deeper insight into the generated embeddings or validating a particular use case. In this work we carried out a comparison of different methods of KGE creation with unsupervised and supervised machine learning tasks. We first generated KGEs from multiple ontologies and biological knowledge bases, and we implemented multiple state-of-the-art methods, and two novel algorithms. Subsequently, we analyzed the generated embeddings using unsupervised clustering algorithms. Furthermore, we evaluated the performance of the embeddings in a GDAs prediction task. Finally, we used the best performing model to predict potential genes associated to intervertebral disc degeneration (IDD).

## METHODS

### Data sources

To build the KG, we mined different type of biological data from publicly available repositories:

#### Protein - protein interactions

We partially integrated data from multiscale interactome (downloaded 29/06/2022) (9). Specifically, the data were integrated from:

- The biological general repository for interaction dataset (BioGRID) (10). This is a repository of manually curated both physical and genetic interactions between proteins from 71, 713 high - throughput and low - throughput publications.
- The database of interacting proteins (DIP)(11) in which only physical protein - protein interactions are reported with experimental and curated evidence.
- Four protein-protein interaction networks coming from verified from the human reference protein interactome mapping project (12)): (HI-I-05: 2, 611 interactions between 1, 522 proteins; HI-II-14 13, 426 interactions between 4, 228 proteins, Venkatesan-09: 233 interactions between 229 proteins; Yu-11 1, 126 interactions between 1, 126 proteins)
- Physical protein-protein interaction from Menche et al. (13)). This repository integrates different resources of physical protein - protein interaction data from experimental evidence. It integrates regulatory interactions from TRANSFAC (14) database, binary interactions from yeast-two-hybrid datasets and curated interactions from IntAct (15), BioGRID and HPRD (16). It integrates also metabolic-enzyme interactions from KEGG (17) and BIGG (18), protein complex interactions from CORUM (19), kinase-substrate interactions from PhosphositePlus (20) and signaling interactions from Vinayagam et al. (21)

Only human proteins for which existed direct experimental evidence of a physical interaction were considered.

#### Ontologies

Ontologies are computational structures that aim to describe and classify the entities belonging to a certain domain in a structured and machine-readable format in order to be implemented in a broad range of applications. The main components of the ontology are classes that represent specific entities and usually are associated with an identifier. These classes are arranged in a hierarchical way from general to more specific and are connected to each other through relations. Finally, ontologies can have also other elements such as metadata, formats and axioms (22) For our purpose we integrated different types of ontologies:

- Gene Ontology (GO) (23), (downloaded 18/07/2022) is a knowledge base that aims to computationally describe biological systems ranging from molecules to organisms, as of 2023 it comprises 43, 248 terms, 7, 503, 460 annotations across 5, 267 species.
- Disease Ontology (DO) (24), (downloaded 02/08/2022) is an ontological structure of standardized disease descriptors across multiple resources. The aim of the project is to provide a computable structure of integrated biomedical data in order to improve the knowledge on human diseases.
- Human Phenotype Ontology (HPO) (25), (downloaded 22/08/2022) is a comprehensive logical structure that describes phenotypic abnormalities found in human diseases. This enables computational inference and interoperability in digital medicine.

We integrated HPO and DO and mapped all the common codes to UMLS CUIS (26) and merged them with GO.

#### Gene products annotations to biological processes

Proteins in the KG were mapped to their specific biological process through GO. GO annotations are statements about the function of a particular gene product, in this way it is possible to obtain a snapshot of the current biological knowledge. We included gene annotations from the gene ontology association file (downloaded 29/06/2022).

#### Gene products annotations to phenotypes

We integrated data of genes associated to phenotypes from 2 sources:

- DisGeNET (27) is one of the largest publicly available collections of genes and variants associated with human diseases. For our purposes we exploited DisGeNET curated (version 7.0) that integrates expert curated human gene disease associations from different data sources DisGeNET integrates GDAs data from curated resources with data automatically mined from the scientific literature using text-mining approaches. To create a dataset, we used curated data from DisGeNET, comprising a total of 84, 037 associations (hereafter considered as positives). We generated the same number of gene-disease non-associations (i.e. negatives) by considering that such associations were not reported in the text – mining version of DisGeNET, hence taking randomly any gene-disease pair not reported as positive.
- HPO gene annotations to phenotypes: HPO (downloaded 02/08/2022) provides a file that links between genes and HPO terms. If variants in a specific gene are associated with a disease, then all the phenotypes related to that specific disease are assigned to that gene.

#### Phenotypes annotated to diseases

We integrated annotations of phenotypes to disease from the phenotype.hpoa file from HPO ontology (downloaded 15/12/2022).

#### Drug-disease associations

We integrated data of drug-disease pairs from the multiscale interactome. This dataset is integrated by a collection of FDA approved treatments for diseases including different sources:

- The drug repurposing database (28) is a database of gold-standard drug-disease pairs extracted from DrugCentral (29) and ClinicalTrials.gov
- The drug repurposing hub (30) is a collection of drug-disease including 4, 707 compounds. The database contains information mined from publicly and proprietary datasets that undergo manual curation.
- The drug indication database (31) integrates data from 12 openly available, commercially available and proprietary information sources.

The dataset was filtered by keeping only human proteins resulting in a total number of drug - disease pairs of 5, 926.

#### Drug - target interaction

We obtained a dataset of drugs and their mode of actions on target proteins by integrating DrugBank (32) and the drug repurposing hub. Proteins that were not included in the protein – protein interaction network were removed.

### Algorithms to create embeddings

Currently we have compared various embedding strategies. We have tested some state-of-the-art algorithms based on different principles and we have implemented two novel methods to generate embeddings, referred as BioKg2vec and DLemb. For all experiments, the embeddings vector dimension was 100.

#### TransE

TransE (33) is an algorithm that relies on a translational - based model. It represents relationships as translations in the embedding space. The principle lays on the assumption that given the triple *“(h, r, t)”*, where h is the head, r is the relation, t is the tail e.g. *“(protein1, interacts with, protein2)”*, the embedding of the tail should be similar to the head embedding plus the relationship embedding.

#### Node2Vec

Node2Vec (34) is an algorithmic framework that learns continuous feature representations for nodes in networks. It relies on the implementation of a biased random walker for exploring the topology of the network. Subsequently, the walks are fed to Word2Vec (35) algorithm to transform entities into numerical representations. We used the implementation available at Stanford Network Analysis Platform (36).

#### Distmult

Distmult (37) is defined as a multiplicative learning framework. It uses an entry-wise product to measure compatibility between head and tail entities. We use the implementation available in pykeen.

#### Graph Convolutional Neural Network (GCN)

GCN (38) applies convolutions to graph data to learn functions on graph features. This approach is inspired from convolutional neural networks (CNN) that uses a function as a filter applied by convolution on the data to learn complex patterns from images. In GCNs these convolutions are applied to spectral representations of the graph with the goal of learning the function. The forward propagation algorithm of the GCN is:

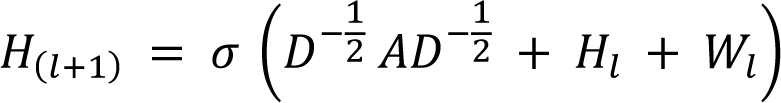

Where 𝜎𝜎 is an activation function such as RELU, 𝐴 = 𝐴 + 𝐼_𝑁_ is the adjacency matrix that encodes the connections in the graph, 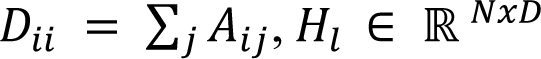 is the matrix of activations in the 𝑙^𝑡ℎ^ layer and 𝑊_𝑙_ is the weight matrix of layer 𝑙. We used the implementation of GCN available in Pytorch Geometric (39).

*BioKg2Vec* BioKg2Vec relies on a biased random-walk approach in which the user can prioritize specific connections by assigning a weight to edges. In the KG defined in this work we used 4 different node-types: drug, protein, function and disease. Then, the probability of visiting a specific neighbour at every step is given by the equation:

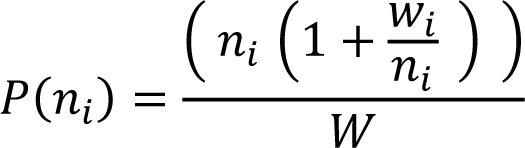

where 𝑃(𝑛_𝑖_) is the probability for the RW to visit a specific node type, 𝑛_𝑖_ is the number of paths leading to the node (of the same type), 𝑤_𝑖_ is the assigned weight (also specific for the type) and 𝑊 equals to the node degree plus the sum of all weights (i.e. ∑_𝑖_ 𝑤_𝑖_) . To detect the optimal weights for the prediction of GDAs we performed a grid search assigning weights prioritizing drug -> protein -> function -> disease. Moreover, the walker stores the information of the visited edge type, and this information is used as input for Word2Vec algorithm in the embedding generation step. Thus, the algorithm handles different edges and nodes behaving differently for each node type being visited and storing the edge type of information too. BioKg2Vec is available at https://zenodo.org/badge/latestdoi/624339823.

#### Dlemb

DLemb is a shallow neural network (NN) that consists of 3 layers: the input layer, embedding layer and output layer. The input layer takes as input KG entities as numbers and outputs them to the embedding layer. Subsequently, embeddings are normalized, and a dot product is calculated between them resulting in the output layer. The model is trained by providing a batch of correct links and wrong links in the KG to provide with positive and negative examples in what can be conceived as a link-prediction task. Embeddings are then optimized for every epoch by minimizing RMSE and using Adam optimization. DLemb is available at https://zenodo.org/badge/latestdoi/635382680.

### Methods to combine embeddings

We used 4 strategies to combine genes and diseases embeddings to obtain GDAs representations: 1) Sum, which consisted on the addition of both vectors; 2) Average, in which we averaged them; 3) concatenation, in which the result is a vector in a larger dimension, representing a pair gene-disease by concatenating both vectors; 4) Hadamard product (i.e. each element is produced by the product of the elements of the two vectors). For this work we produced embeddings of fixed dimension (i.e. 100) in the space of reals (i.e.ℝ^100^).

### Principal component analysis (PCA) and unsupervised clustering of embeddings

We performed a PCA on the embeddings to reduce the dimensionality of the 100 vectors. Then we plotted the first 2 dimensions of the embeddings related to genes and diseases and visualized how they could cluster different classes using the k-means algorithm. Specifically, we used function and compartment-based classification to group gene products in different categories: secreted, transcription factors and transport (40). For diseases, we used annotations from UMLS to ICD-9 (41), that classifies diseases in 17 macro classes including mental disorders, congenital anomalies, and infectious and bacterial diseases. We then used various evaluation scores for the comparison, such as the silhouette score, defined as:

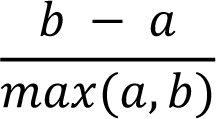

where 𝑏 is the mean distance between a sample and all other points in the nearest cluster (nearest – cluster distance) and 𝑎𝑎 is the mean distance between a sample and all other points in the same class (inter – cluster distance). We calculated this score for different cluster sizes ranging from 1 to 10 for genes (the gold standard number of clusters is 3) and from 10 to 20 for diseases (the gold standard number of clusters is 15).

Finally, we evaluate the homogeneity score, defined as:

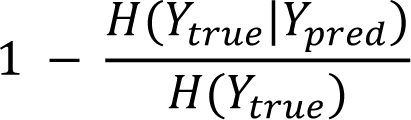

That is a measure that quantifies the similarity of samples in each cluster. Where the 𝑌_𝑡_ is the number of classes, 𝑌_*pred*_ is the number of clusters and term 𝐻(𝑌_*true*_|*Y_pred_*) which represents the ratio between the number of classes 𝑌_𝑡_ in cluster 𝑌_*pred*_ and the total number of samples in cluster 𝑌_*pred*_. When all the entities in the cluster belong to a class the homogeneity score equals 1.

### Grid-Search to select the best predictive model

We performed a grid search cross-validation to find the best combination of embedding creation algorithm, GDAs representation and predictive ML and DL algorithms implemented in Scikit-learn (42) and Pytorch (43) respectively. In the grid-search experiment we created a KG in which we integrated all the biological data and 80% of curated GDAs from DisGeNET. We tested the predictions in the remaining 20% of GDAs. With this, we tested a total of 120 combinations for the grid-search. For each algorithm we fitted a grid of parameters maximizing the area under the receiver operating-characteristic curve (AUROC) table 1. Then the best parameter combination was evaluated on the test set by assessing additional evaluation metrics, such as:

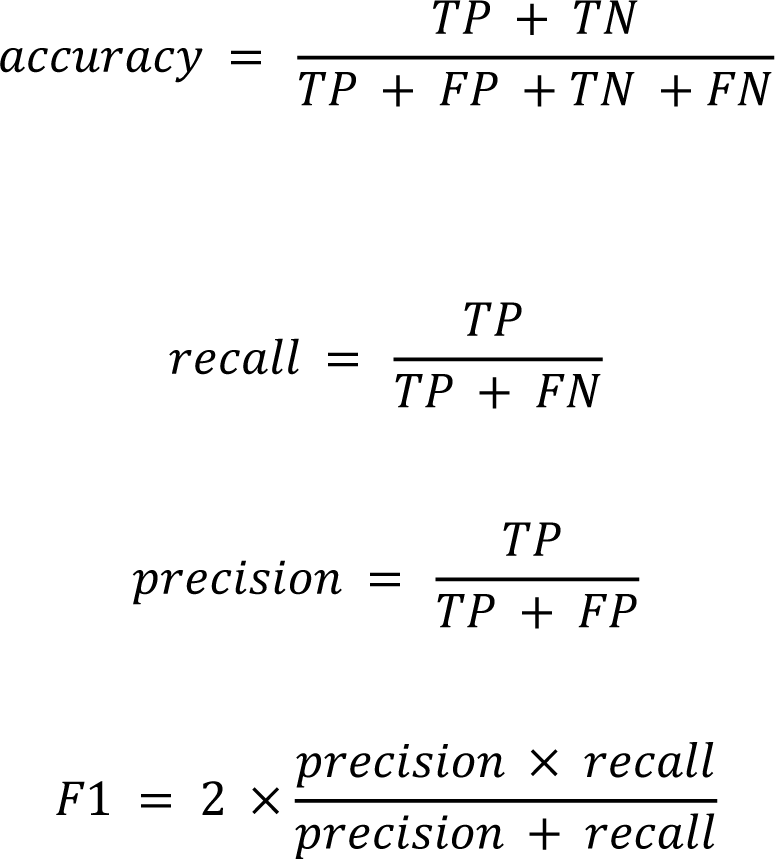

**Table 1:**
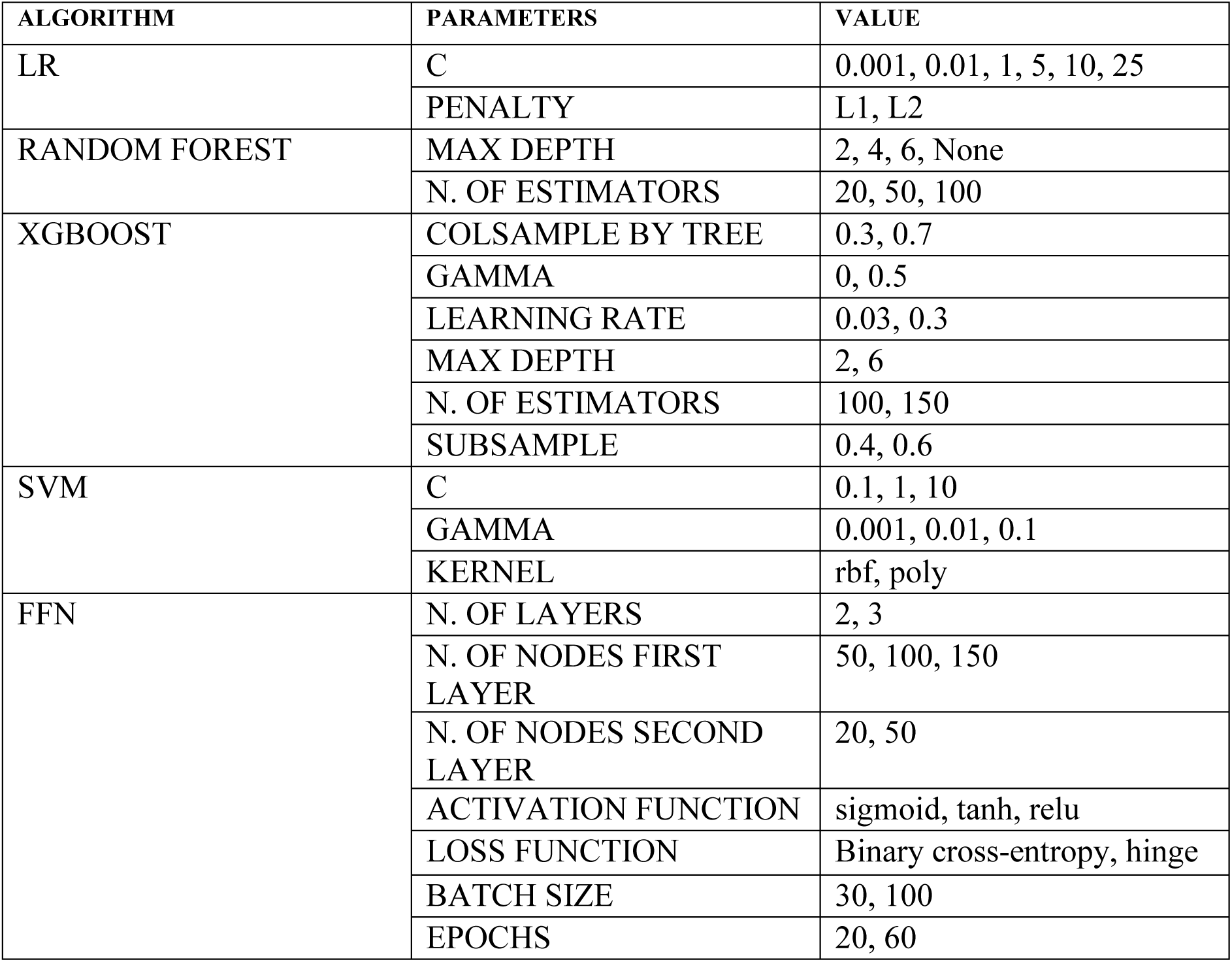
Search spaces of the algorithms tested during the grid search cross validation.

Plus AUROC, that had already been used for the grid-search optimization and Area under the precision recall curve (AUPRC). AUROC is the amount of area covered under the curve of the recall versus the False Positive Rate (FPR) defined as:

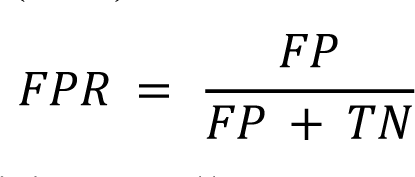

and AUPRC the area under the precision-recall curve.

TP is the number of positives predicted correctly and FP those erroneously predicted as negatives, while FN and TN are those negatives erroneously predicted positives or correctly predicted negatives, respectively.

### Ontology preprocessing and heterogeneous data integration

Once we selected the model with the highest predictive power, we investigated the influence of integrating heterogeneous biological data in the KG on the GDAs predictions. For this experiment we only used ontological data. Ontologies are complex, standardized data structures composed of classes, relations, axioms and metadata all of which are included in the raw ontology. Moreover, we tested the effect of implementing a pre-processing step in the ontology in which only classes and relations were maintained as a graph structure (axioms and metadata were excluded). We studied the following combinations of data sources:

1. HPO + HPO annotations raw
2. HPO + HPO annotations pre-processed
3. HPO + HPO annotations + GO + GO annotations (all) pre-processed

We used a comparison based on two metrics. For this experiment, we created embeddings with the DLemb algorithm, using concatenation for GDAs representation, and SVM as classification algorithm. For the processing of the ontologies nxontology and pronto (44) python libraries were used.

### Influence of GDAs in the KG for GDA-predictions

We tested the influence of adding increasing GDAs proportions in the KG. For this experiment, we used 0% 20% 50% and 100% of DisGeNET and we included it in the KG. Then we generated embeddings from the KGs with DLemb and we trained a SVM on 80% of DisGeNET. We tested the model on the 20% of remaining associations and calculated AUROC and AUPRC as evaluation metrics.

### Comparison with random generated embeddings

In order to show that the information is efficiently translated from the KG to the vectorial space we compared the performance of DLemb generated embeddings and random embeddings of the same size. We wanted to test if embeddings could describe diseases and genes and thus predict gene-disease associations merely due to their construction using KGs. To test this, we generated 100-dimension random embeddings for every gene and disease and represented GDAs with concatenation. The number of associations is a latent variable that can be learned by ML producing good predictions. This can be considered a potential bias. Therefore, we further tested the effect of removing the number of associations by stratifying DisGeNET data. We obtained 23 classes in which the number of associations for every disease have a maximum difference of 20. Then we selected a disease belonging to every class, generated negative associations and performed a five-fold cross validation on the data with the best performing algorithm. We evaluated Accuracy, precision, recall, f1 score and AUROC across every fold.

### Generalizability of the model

The predictive model selected was tested to predict associations for diseases not used in the training set. The rationale behind this experiment was to understand the capabilities of the model to predict gene-disease associations of new diseases, proving that the biological information encoded in the embeddings was generalizable.

To assess this, we trained the model on GDAs belonging to diseases of a specific ICD9 disease class and then we tested the model on all other classes.

### Intervertebral Disc Degeneration Biomarker Prediction

We tested the model to predict genes associated to IDD. We used the model selected trough grid search cross validation with concatenation of the embeddings for the GDAs representation. Lastly, we performed a function enrichment analysis using g:Profiler (45) on the set of prioritized genes with probability higher than 0.95 to be associated to IDD.

## RESULTS

### Knowledge Graphs

We integrated multiple sources of data in the form of KG for a total of 103, 206 nodes and 2, 208, 829 edges. The KG contains 4 types of nodes: drugs (n = 3, 101), phenotypes (n = 35, 431), proteins (n = 21, 106) and functions (n = 43, 568). These entities are connected by 81 different types of relationships represented as edges. The relationships are obtained through different data sources, 17, 660 proteins interacting among each other (775, 252 edges), 19, 409 proteins annotated to 18, 813 biological functions (303, 404 edges) and 9, 965 proteins annotated to 18, 681 phenotypes (29, 6182 edges). Moreover, drug information was included: 1, 661 drugs annotated to 840 phenotypes for a total of 5, 926 edges and 2, 887 connected to 2, 074 proteins they target for a total of 14, 491 edges. The degree distribution of the graph follows a scale free law (Supplementary figure 1) (46).

### Unsupervised clustering of the embeddings reflects biological classification

From the KG, we generated embeddings using six algorithms. Figure 1 shows the two principal components of gene and disease embeddings. The embeddings tend to differentiate among gene products belonging to different groups: secreted, transcription factors, and transporters (Figure 1 A to G). Regarding the algorithms, TransE, BioKG2vec, and DLemb can distinguish better the genes than the rest of algorithms. In Figure 1, G to L only 3 categories of diseases are represented, corresponding to the ICD chapters congenital anomalies, infectious and parasitic diseases, and mental disorders. As above, algorithms TransE, BioKG2Vec and DLemb visually distinguished disease classes better than others.

**Figure 1:**
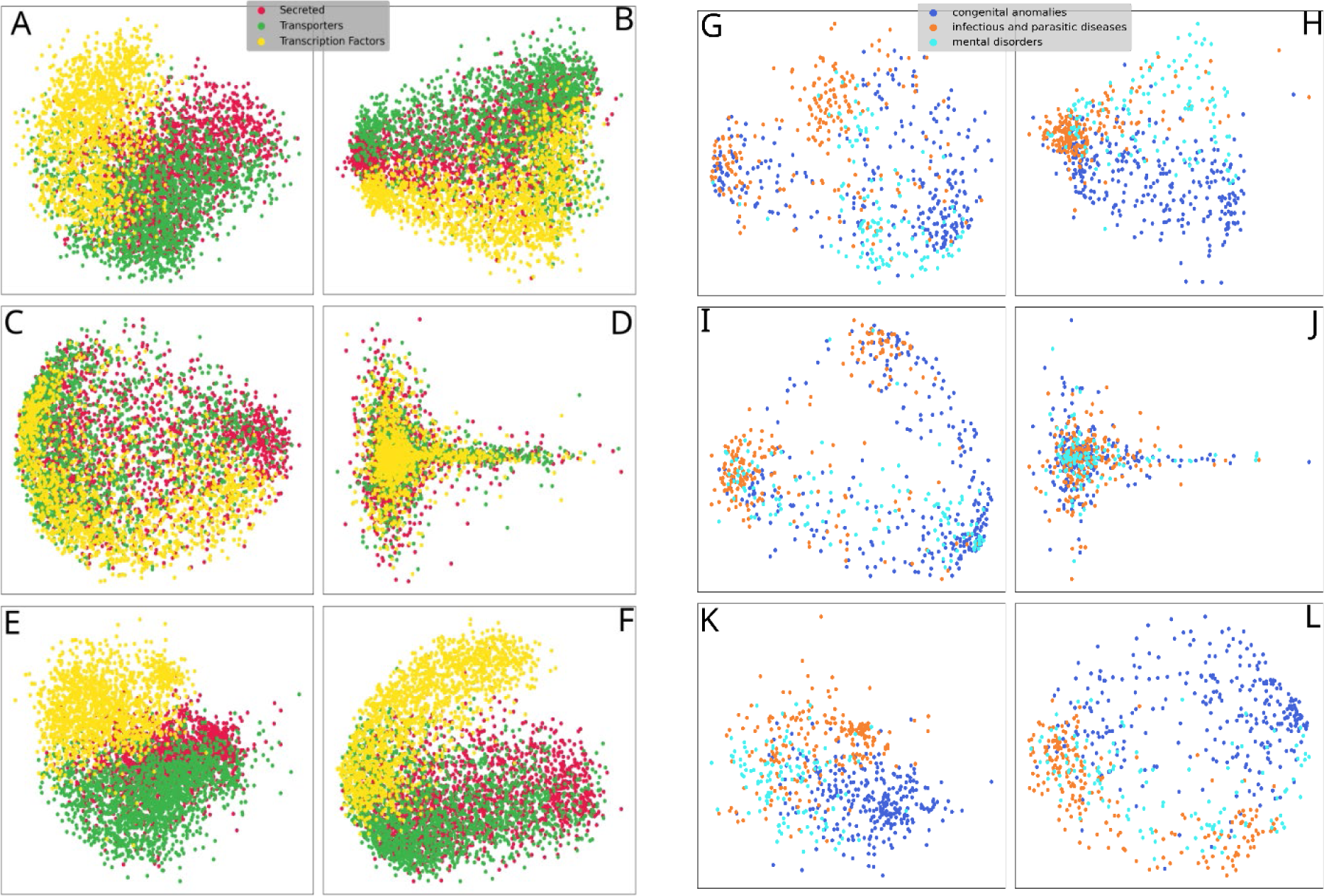
PCA of gene-embeddings (panels A to F) and disease-embeddings (panel G to L) generated with TransE (A), Node2Vec (B), DistMult (C), GCN (D), BioKG2Vec (E) and DLemb (F).

The algorithm producing the best clustering of disease classes and gene products was BioKG2Vec, which has a higher homogeneity score for both, genes and diseases. Furthermore, these embeddings also reach the highest silhouette score when the number of clusters = 3 i.e. the number of gene product classes (Supplementary figure 2). For the case of diseases, the silhouette score of the embeddings produced with any algorithm couldn’t match the gold standard number of clusters (Supplementary figure 3).

The embeddings of gene products with BioKG2Vec produced homogeneous clusters (Supplementary figure 4). Similarly, disease embeddings generated with the same algorithm cluster diseases of the same class, with the exception of few classes (such as those of pathological sign and symptoms class, both, distributed across many clusters) and of congenital anomalies, that can be found in at least two clusters (Supplementary figure 5).

### Model selection through grid search cross validation

The best performing combination for GDA prediction was DLemb deep learning model for creating embeddings coupled with concatenation of the gene and disease embedding as association representation and SVM with parameters C = 10 and kernel = rbf as classification algorithm. The whole output of the experiment is available in Supplementary table 1. The following experiments were run using this combination.

### The pre-processing of the ontology affects prediction of GDAs

Pre-processing the ontologies leads to better AUROC and AUPRC compared to using embeddings generated with raw data. Nevertheless, adding heterogeneous data in the KG did not significantly affect the predictions of GDAs (Table 3). Integrating more data leads to similar performances which can be appreciated when comparing the results of generating the KG using HP data with HP and GO data.

### The amount of training GDAs in the KG affects the prediction of GDAs

We tested the effect on the predictions caused by the increase of GDAs in the KG. We expect that increasing the amount of GDAs in the KG will increase the quality of the predictions. Figure 2 shows that the increase in the number of GDAs used for training the knowledge graph embeddings increases the values of AUROC and AUPRC.

**Figure 2:**
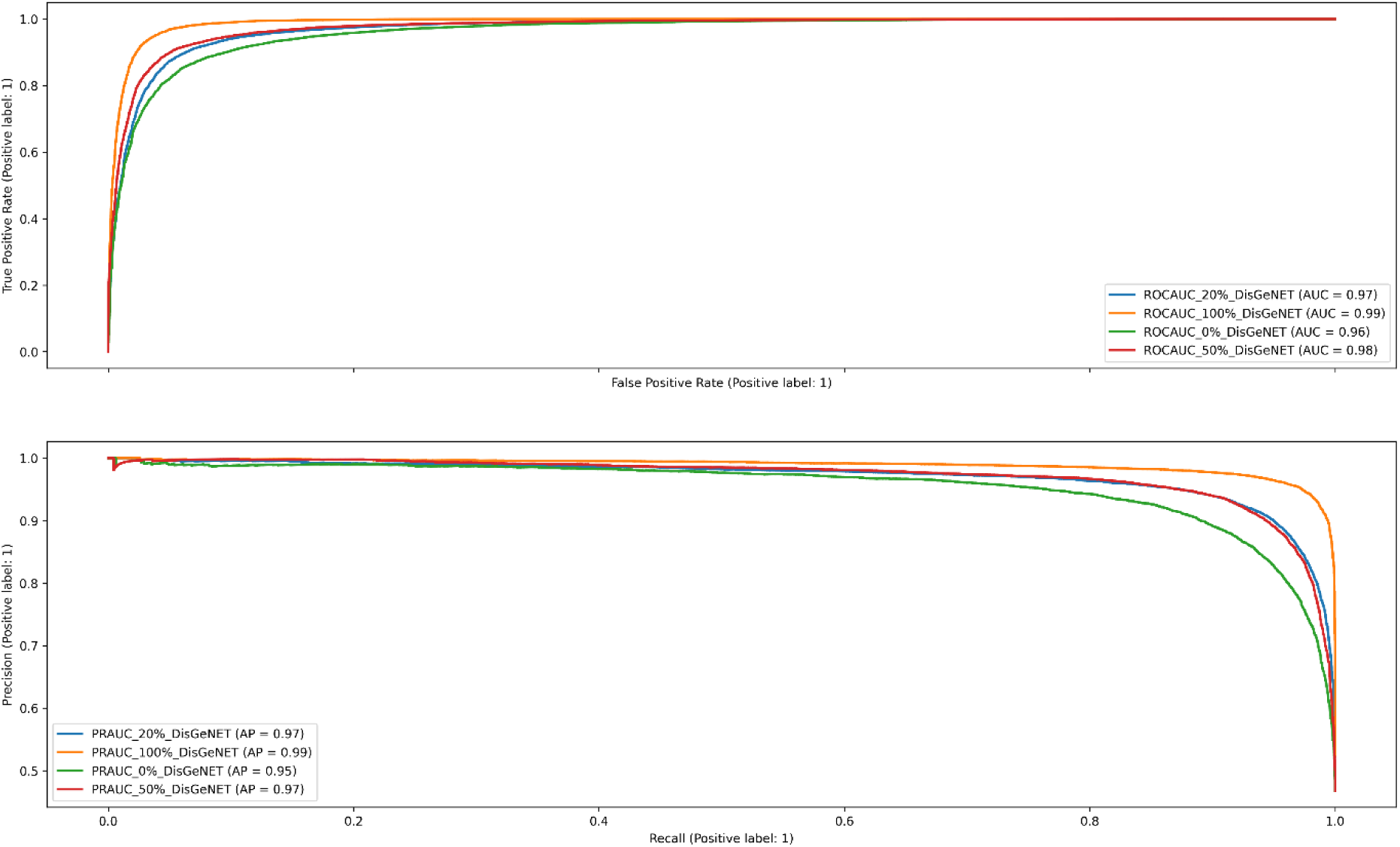
ROC and precision-recall curves of the prediction of GDAs. Several KG embeddings are obtained using increasing percentages of known GDAs from 0 to 100%. Note: embeddings were generated with DLemb, using concatenation for combining embeddings and SVM for the classification/prediction algorithm.

### Comparison with randomly generated embeddings

In Supplementary table 2 are showed the result of the experiment relative to the GDA prediction relative to diseases with different number of associations in DisGeNET. DLemb embeddings reach an average AUROC of 0.88 while random generated embeddings have random metrics. These results are due to the biological information intrinsic to the embeddings since the effect of the number of GDAs was prevented by selecting associations of one disease only. In fact, the number of associations is a latent that is learned by the model.

### Model generalization across different disease classes

Figure 3 shows the performance of the model trained on a specific ICD9 disease class and tested on all the others. The performance on training and testing on the same class leads to high performance. However, the biological information encoded in the embeddings is translated across disease classes and we can see that some pairs of disease classes achieved a noteworthy prediction (e.g the model trained for neoplasms predicts genes associated with diseases of circulatory system with an AUROC > 0.7. As expected, random generated embeddings show AUROC in the heatmap with random values. Thus, the lack of biological information encoded in the embeddings implies the incapability of generalizing predictive models across disease classes.

**Figure 3:**
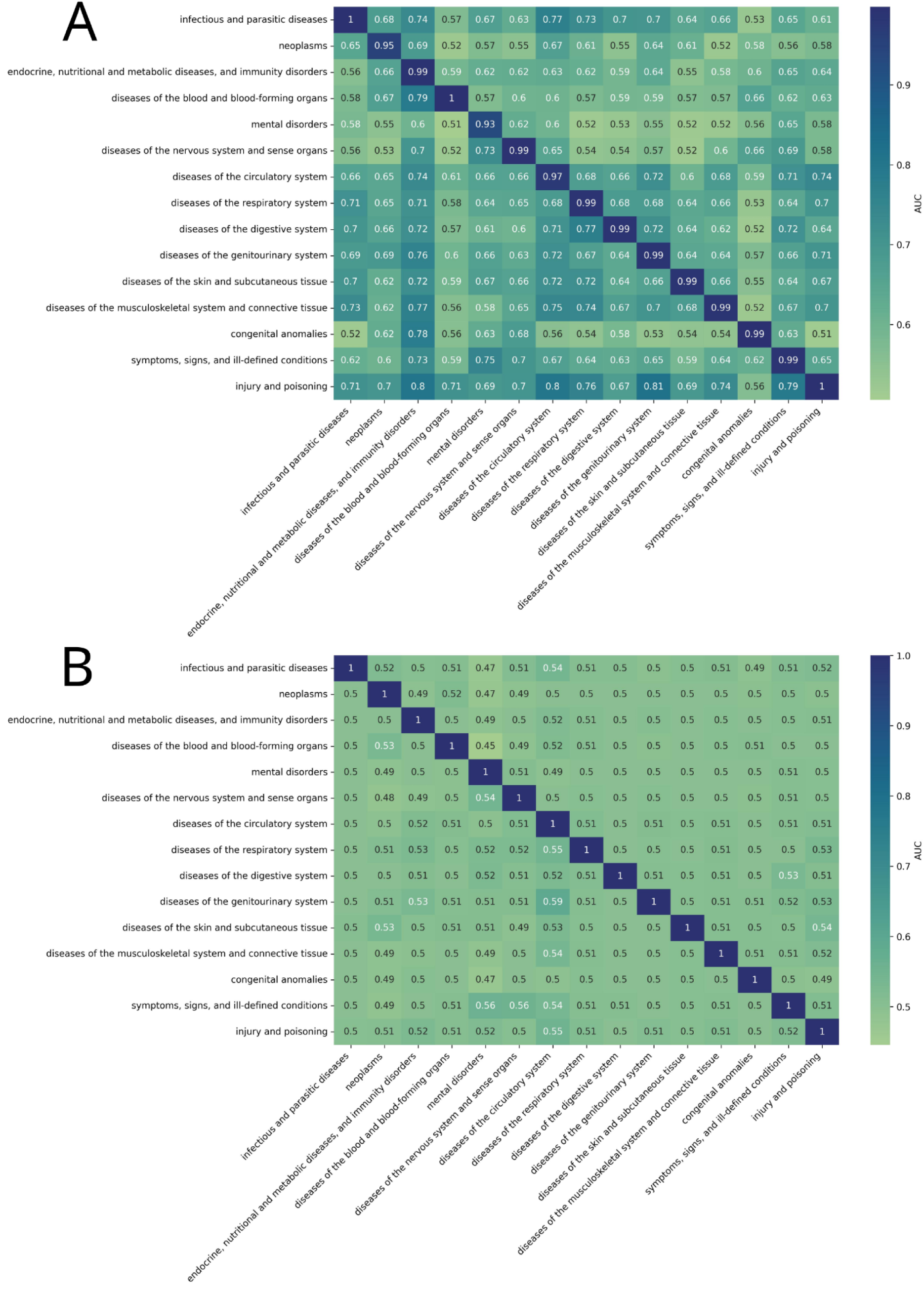
AUROC of the best performing combination for the prediction on different disease classes. A) The embeddings were generated with DLemb deep learning model, the GDAs representation was concatenation, and the algorithm was SVM with parameters C = 10 and kernel = rbf. We show the results of training a model on a specific disease class (shown in the rows), and then testing on the others (columns). B) Results for randomly generated embeddings, the AUROC shows random values.

### Biomarkers prediction for intervertebral disc degeneration

Finally, we used the selected prediction model with best parametrization to predict GDAs for IDD. IDD is one of the main causes of low back pain (LBP), the largest cause of morbidity worldwide affecting 80% of people from Western countries during their lifetime (47). IDD consists of the gradual deterioration of the intervertebral disc (IVD) in which the content of collagen and glycosaminoglycan decreases, and it becomes more dehydrated and fibrotic. Due to this, its anatomical areas nucleus pulposus (NP) and anulus fibrosus (AF) becomes less distinguishable (48). Also, during IDD there is a catabolic shift in the biochemical processes of the disc environment with an increased expression of matrix degrading enzymes promoted by catabolic cytokines and vascularization of the tissues (49). According to DisGeNET (curated sources), IDD is associated to TGFβ-1, HTRA1 and SPARC. We ran predictions for 21, 078 genes by using concatenation as a GDAs representation, of those 364 were predicted to be associated to the disease and 68 with a probability > 0.95. The results of the top 10 prioritized genes are showed in Table 2.

**Table 2:** Top 10 genes prioritized from the best performing model

|  | Homogeneity score |  |
| --- | --- | --- |
|  | Genes | Diseases |
| DLemb | 0.3 | 0.2 |
| DistMult | 0.02 | 0.11 |
| GCN | 0.002 | 0.02 |
| BioKG2Vec | **0.7** | **0.3** |
| Node2Vec | 0.03 | 0.14 |
| Trans-E | 0.4 | 0.2 |

**Table 3:** AUROC and AUPRC of different experiments of GDAs predictions using human phenotype ontology (HPO) + annotations (A), HPO ontology processed (B) and HPO + Gene Ontology (GO) + GO annotations. The embeddings were generated with DLemb and we used SVM as predictive algorithm, and operator concatenation for combining the embeddings.

| Experiment | AUROC | AUPRC |
| --- | --- | --- |
| HPO + HPO annotations raw | 0.77 | 0.88 |
| HPO + HPO annotations processed | **0.95** | **0.98** |
| HPO + HPO annotations + GO + GO annotations processed | 0.94 | 0.97 |

Interestingly, none of the resulting prioritized genes had previously been associated with IDD in DisGeNET (curated dataset). TGFβ2, COL2A1, MMP2, MMP9 have widely been associated with IDD in the scientific literature according to the same database. EXT1 and EXT2 are enzymes responsible of synthesis of heparan sulphate disaccharide chain an important component of glycosaminoglycans in the IDD (50). It was shown that mouse with knockdown EXT1 gene showed a deranged intervertebral disc and joint formation (51). An important part of IDD is the vascularization of the IVD, in fact during degeneration we assist to an ingrowth of blood vessels and innervation of the tissue. This process is driven by VEGF and its receptors (one of which is KDR) (52) and it has been associated with back pain (53). SMAD3 is an important gene for the homeostasis of the disc having a role in the TGF-β pathway (54). Finally, SERPIN1A is a serine protease inhibitor which overexpression in NP cells was showed to promote cell viability and inhibited cell apoptosis and senescence and regulation of IDD grade in rat models (55). These genes were shown to have a role in IDD and could be further investigated to elucidate the mechanisms that lead to the degeneration of the disc.

To further explore the biological functions of these candidate genes, we performed an enrichment analysis (Figure 4). The top prioritized genes are enriched in processes related to the extracellular matrix organization, pathways related to collagen formation, and extracellular matrix degradation, all of them related to IDD.

**Figure 4:**
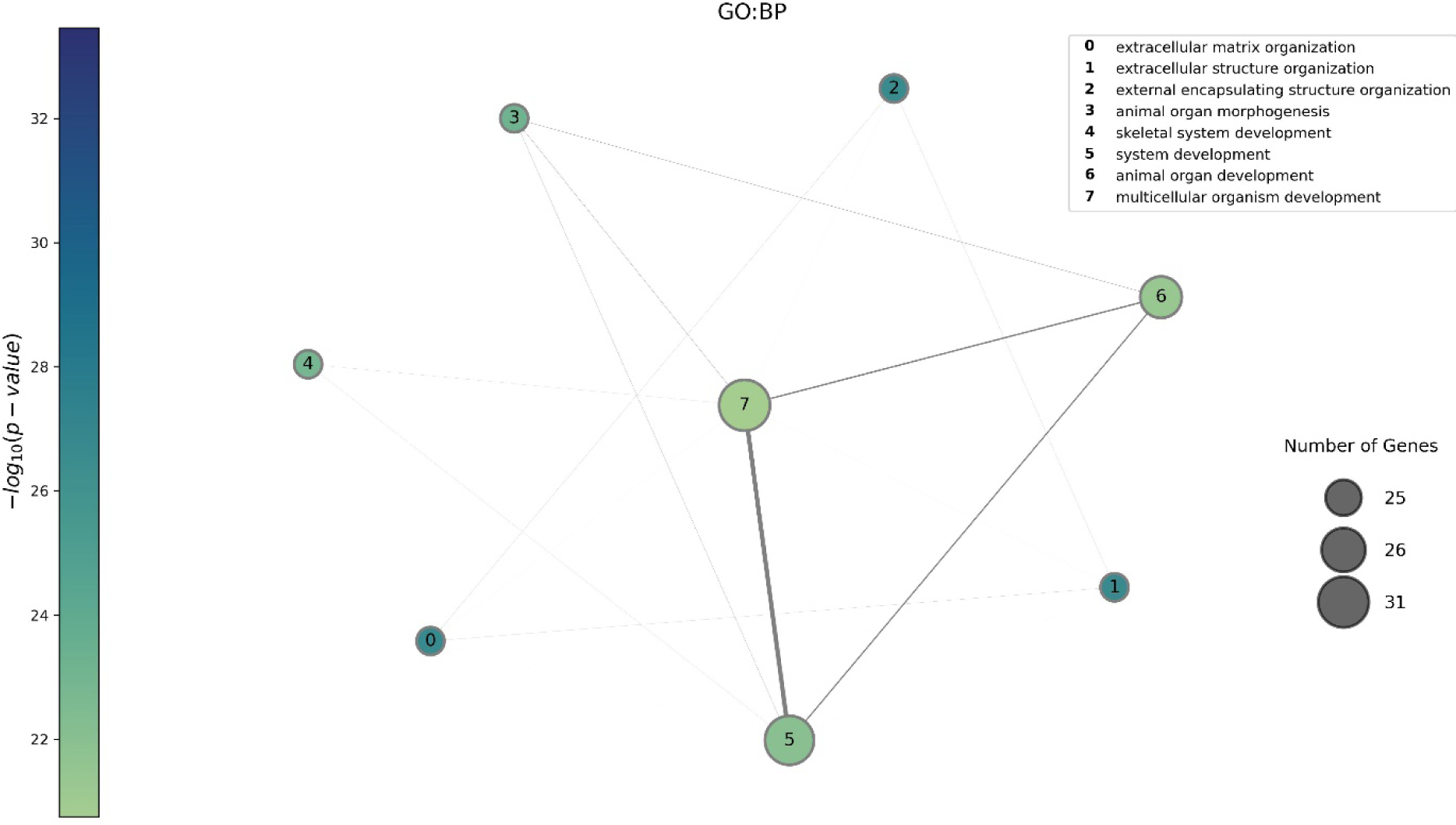
Gene ontology biological processes (GO:BP) function enrichment analysis on the genes with probability higher than 0.95 to be associated to C0158266 (n=68). To run the functional enrichment, we used g:Profiler. The nodes correspond to the pathway enriched in the gene set, their size is proportional to the number of genes belonging to that specific pathway and the colour is related to the significance of the enrichment in the gene set (calculated through hypergeometric distribution). An edge exists between 2 nodes if there are genes shared between the two pathways and the width of the edge is proportional to the number of the genes shared.

**Table 2:** Homogeneity score of K – means algorithm calculated for genes (number of clusters = 3) and diseases (number of clusters = 16). True labels are classification from ICD-9 and HPA for diseases and genes respectively

| Gene ID | Gene Symbol | Probability |
| --- | --- | --- |
| 4318 | MMP9 | 1 |
| 2131 | EXT1 | 1 |
| 7042 | TGFB2 | 1 |
| 4313 | MMP2 | 1 |
| 5176 | SERPINF1 | 1 |
| 1280 | COL2A1 | 1 |
| 7422 | VEGFA | 1 |
| 4088 | SMAD3 | 0.996 |
| 3791 | KDR | 0.995 |
| 2132 | EXT2 | 0.995 |

## DISCUSSION

In this work we investigated how KG embeddings perform to predict gene-disease associations. First, we generated several KGs by implementing heterogeneous biological information such as protein-protein interactions, gene-disease associations, drug-disease associations and drug-protein interactions, and ontologies The integration of multiple knowledge-based datasets prevented us to use syntactic-based approaches for embedding-creation such as OPA2VEC (56). Syntactic approaches rely in the set of axioms only for obtaining the embeddings without the intermediate graph-based representation [57], so the input of the algorithm must be in Web-Ontology Language (OWL) format. Moreover, the integration of different ontologies is a challenging task and an active research topic (58).

Here, we performed a comprehensive evaluation of different methods for KGE creation, and we implemented two new algorithms that outperformed existing methods: BioKG2Vec and DLemb. BioKG2Vec is a biased random walk approach that explores the network by prioritizing certain node classes with starting weights defined by the user. In this way, the walker is not merely exploring network topology but is behaving differently based on the node-type. Also, edge-features are introduced in the walk and fed to Word2Vec. These features make the novelty of this approach, achieving the highest homogeneity score in separating disease and gene classes. DLemb is a shallow NN composed by 3 layers. Despite its simplicity, the embeddings created through this model reached the highest AUROC in the prediction of GDAs according to a grid-search cross validation experiment.

We performed an extensive analysis of the embeddings with unsupervised machine learning methods: we tested the effect of integrating different types of data (ontologies and GDAs), we also compared the prediction of GDAs with the use of random features. In the analysis we showed that increasing proportions of GDAs information in the KG improves the performance of the model. This suggests that task specific embeddings and data improves the predictions, probably because it learns the relevant features as it was shown somewhere else(59). Finally, we predicted new genes associated to IDD, showing the use of KG embeddings to infer biomarkers for diseases with a limited genetic knowledge: the training set (DisGeNET curated) had only 3 associations and the model prioritized 364 genes. These 364 genes reflected the biology underlying IDD. Polygenicity is a feature of common diseases that hinders the understanding of the biology underlying the development of complex conditions (60).

We acknowledge several limitations to this study, first the lack of interpretability of the embeddings. Embeddings are created through neural networks that doesn’t allow to understand the features obtained. Second, there is a hidden bias in the data because some genes and diseases have been more studied than others. This implies to handle an unbalanced number of associations. In fact, many diseases are associated with few genes and few diseases are associated with many genesThis feature is learnt in the model, and it is intrinsic to the current biological knowledge. This also reflects the biology underlying complex and rare diseases that are associated with many and few genes respectively.

## CONCLUSIONS

In this work we carried out an extensive investigation on KGE from the generation and evaluation of the produced embeddings to the development of two new models for KGE generation and the utilization of the so created embedding in a GDAs task. We showed that embeddings can effectively be implemented in the biomedical field in order to infer new knowledge over a certain domain. Nevertheless, many challenges remain open that requires interdisciplinary collaboration to reach better outcomes in healthcare sector.

## DATA AVAILABILITY

KGs and embeddings generated with Node2Vec, DLemb and BioKG2Vec, the pre-trained trained model for GDA-predictions, as well as the code to reproduce the analysis are available at https://zenodo.org/badge/latestdoi/655840675.

## SUPPLEMENTARY DATA

Supplementary Data are available at NAR online.

## ACKNOWLEDGEMENTS

BO acknowledges support from MCIN and the AEI (DOI: 10.13039/501100011033) by grants PID2020-113203RB-I00 and “Unidad de Excelencia María de Maeztu” (ref: CEX2018-000792-M)

