## Supplementary Materials for "PREDICTING GENE DISEASE ASSOCIATIONS WITH KNOWLEDGE GRAPH EMBEDDINGS FOR DISEASES WITH CURTAILED INFORMATION"

SUPPLEMENTARY MATERIAL

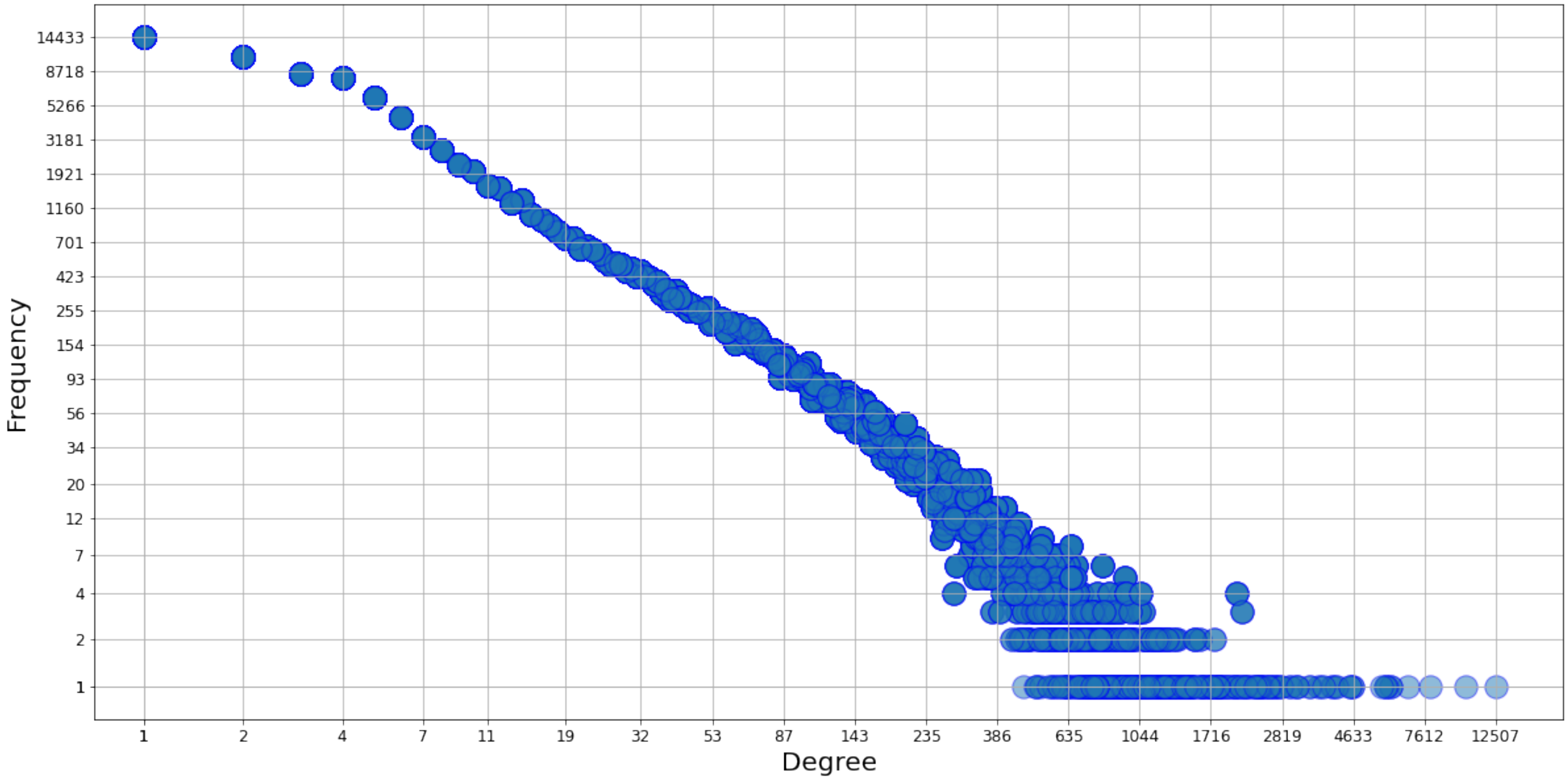


*Supplementary figure 1: Degree distribution of the knowledge graph, x axis represents the degree i.e. number of edges adjacent to a specific node and y axis is the frequency of nodes with that specific degree in the graph. The KG follows a scale free degree distribution*

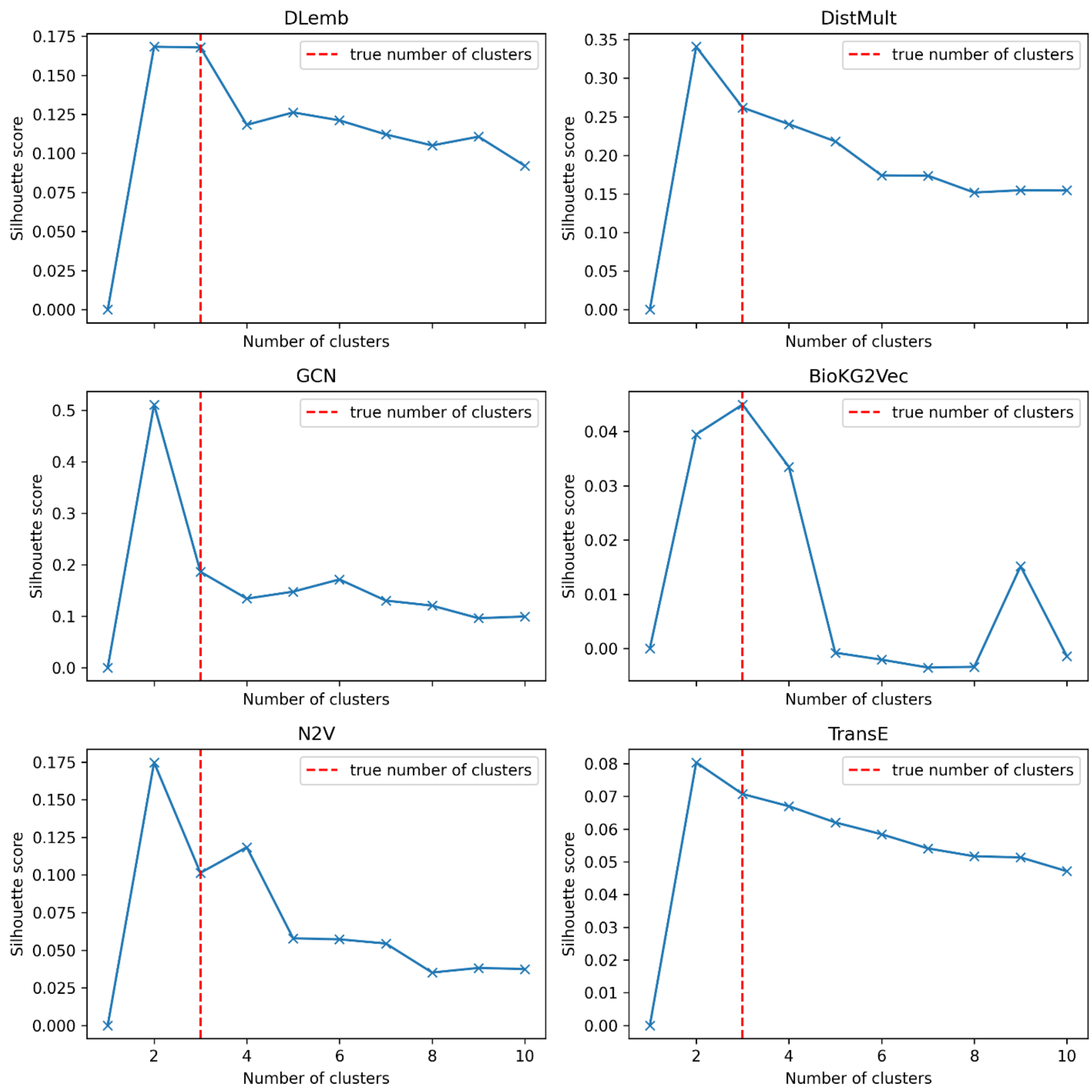


*Supplementary figure 2: Silhouette scores calculated for different numbers of K-means clusters, the red line represents n = 3 i.e. the actual number of gene classes from HPA (Secreted, Transcription Factors, Transporters). BioKG2Vec has the highest silhouette score when the number of clusters = 3.*

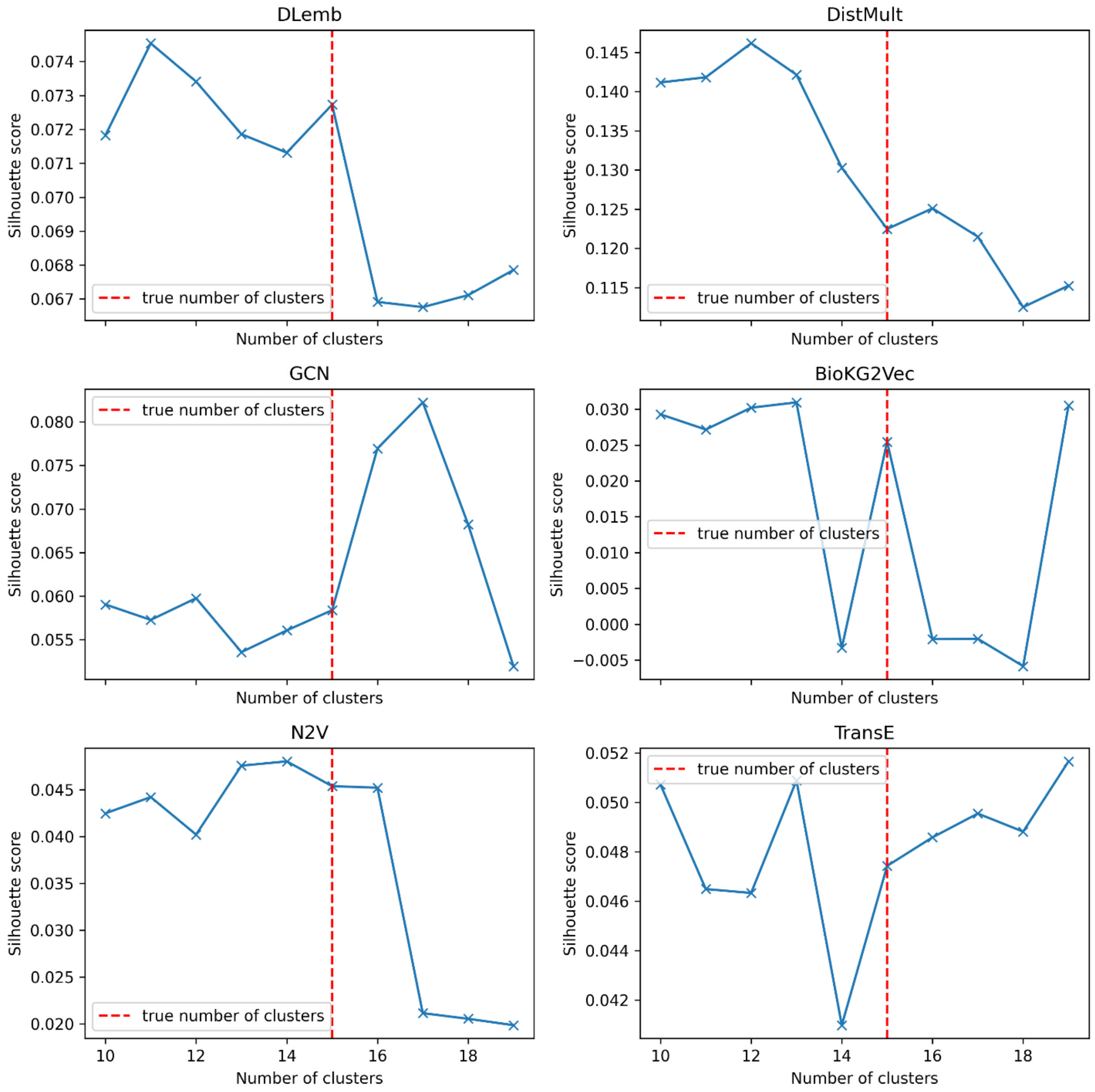


*Supplementary figure 3: Silhouette score calculated for different numbers of K-means clusters, the red line represents n = 16 i.e. the actual number of disease classes from ICD-9.*

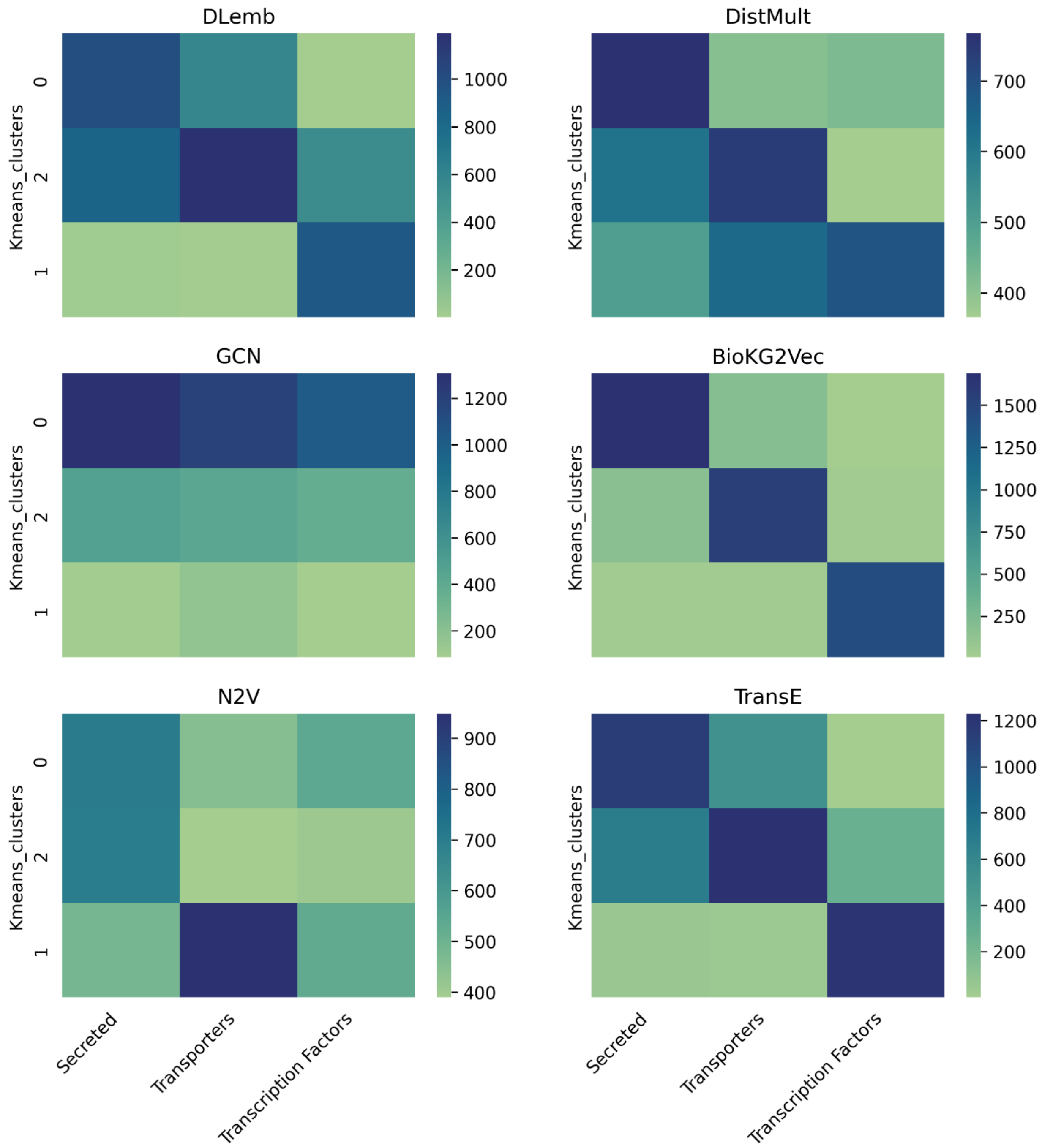


*Supplementary figure 4: K-means clusters on gene product embeddings separated by Human Protein Atlas protein categories. On the y axis are the 3 clusters produced from the algorithm and on the x axis the protein classes. The color indicates the number of gene products in each cluster.*

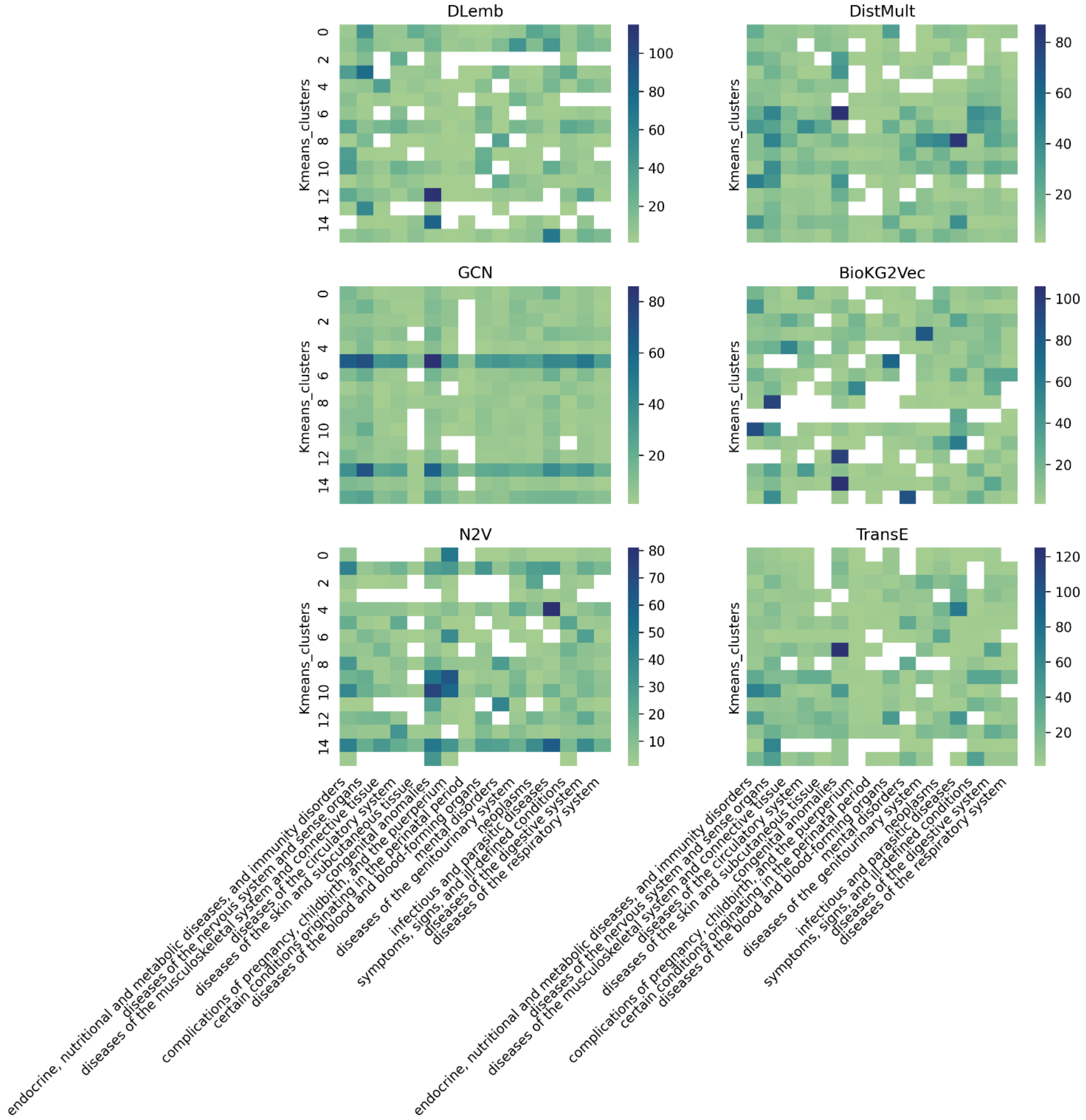


*Supplementary figure 5: K-means clusters on disease embeddings separated by ICD-9 disease codes. On the y axis are the 15 clusters produced from the algorithm and on the x axis the disease classes. The color indicates the number of diseases in each cluster. BioKG2Vec reach highest separation of disease classes*

*Supplementary table 1: Results of grid search cross-validation. The results are ordered by AUC. DLemb and BioKG2Vec with Concatenation GDA representations are the best performing algorithms. Svm is support vector machine, lr logistic regression, xgb xgboost, rf random forest, and ffn feedforward neural network.*

| Grid_Instance | F1 | PRECISION | RECALL | ACCURACY | AUCROC | PRA |
|---|---|---|---|---|---|---|
| DLEMB_Concatenation_svm | 0.95 | 0.97 | 0.95 | 0.95 | 0.95 | 0.95 |
| DLEMB_Concatenation_xgb | 0.94 | 0.97 | 0.94 | 0.94 | 0.94 | 0.95 |
| BioKG2Vec_Concatenation_svm | 0.93 | 0.98 | 0.91 | 0.93 | 0.94 | 0.95 |
| DLEMB_Concatenation_ffn | 0.94 | 0.96 | 0.94 | 0.94 | 0.94 | 0.95 |
| BioKG2Vec_Concatenation_ffn | 0.93 | 0.98 | 0.91 | 0.92 | 0.93 | 0.95 |
| TransE_Concatenation_svm | 0.94 | 0.96 | 0.95 | 0.94 | 0.93 | 0.94 |
| TransE_Concatenation_ffn | 0.93 | 0.97 | 0.92 | 0.93 | 0.93 | 0.94 |
| TransE_Concatenation_xgb | 0.93 | 0.96 | 0.94 | 0.93 | 0.93 | 0.94 |
| DLEMB_Hadmard_svm | 0.93 | 0.96 | 0.93 | 0.93 | 0.93 | 0.94 |
| DLEMB_Hadmard_ffn | 0.93 | 0.96 | 0.93 | 0.93 | 0.93 | 0.94 |
| DLEMB_Hadmard_xgb | 0.93 | 0.97 | 0.92 | 0.93 | 0.93 | 0.94 |
| DLEMB_Average_svm | 0.93 | 0.96 | 0.93 | 0.93 | 0.93 | 0.94 |
| DLEMB_Concatenation_rf | 0.94 | 0.95 | 0.95 | 0.94 | 0.93 | 0.94 |
| DLEMB_Sum_svm | 0.93 | 0.96 | 0.92 | 0.93 | 0.93 | 0.94 |
| DLEMB_Sum_ffn | 0.93 | 0.96 | 0.93 | 0.93 | 0.93 | 0.94 |
| DLEMB_Hadmard_rf | 0.92 | 0.96 | 0.92 | 0.92 | 0.92 | 0.94 |
| BioKG2Vec_Concatenation_xgb | 0.92 | 0.97 | 0.91 | 0.92 | 0.92 | 0.94 |
| BioKG2Vec_Average_svm | 0.91 | 0.98 | 0.88 | 0.91 | 0.92 | 0.94 |
| DLEMB_Sum_xgb | 0.92 | 0.96 | 0.93 | 0.92 | 0.92 | 0.93 |
| DLEMB_Average_xgb | 0.92 | 0.96 | 0.93 | 0.92 | 0.92 | 0.93 |
| DLEMB_Average_ffn | 0.93 | 0.95 | 0.93 | 0.93 | 0.92 | 0.93 |
| DistMult_Concatenation_ffn | 0.92 | 0.96 | 0.92 | 0.92 | 0.92 | 0.93 |
| DLEMB_Hadmard_lr | 0.92 | 0.96 | 0.92 | 0.92 | 0.92 | 0.93 |
| BioKG2Vec_Sum_svm | 0.91 | 0.98 | 0.87 | 0.90 | 0.92 | 0.94 |
| N2V_Concatenation_svm | 0.92 | 0.95 | 0.92 | 0.92 | 0.92 | 0.93 |
| DistMult_Concatenation_xgb | 0.92 | 0.95 | 0.93 | 0.92 | 0.92 | 0.93 |
| BioKG2Vec_Sum_ffn | 0.90 | 0.98 | 0.86 | 0.90 | 0.91 | 0.94 |
| BioKG2Vec_Average_ffn | 0.90 | 0.98 | 0.86 | 0.90 | 0.91 | 0.93 |
| TransE_Hadmard_ffn | 0.91 | 0.96 | 0.90 | 0.91 | 0.91 | 0.93 |
| N2V_Concatenation_ffn | 0.91 | 0.95 | 0.92 | 0.91 | 0.91 | 0.92 |
| DistMult_Average_ffn | 0.92 | 0.94 | 0.93 | 0.92 | 0.91 | 0.92 |
| TransE_Hadmard_xgb | 0.91 | 0.96 | 0.90 | 0.91 | 0.91 | 0.93 |
| DLEMB_Sum_rf | 0.92 | 0.94 | 0.94 | 0.92 | 0.91 | 0.92 |
| TransE_Concatenation_rf | 0.92 | 0.93 | 0.95 | 0.92 | 0.91 | 0.92 |
| DistMult_Hadmard_ffn | 0.91 | 0.95 | 0.91 | 0.91 | 0.91 | 0.92 |
| DLEMB_Average_rf | 0.92 | 0.94 | 0.94 | 0.92 | 0.91 | 0.92 |
| DistMult_Sum_ffn | 0.92 | 0.93 | 0.94 | 0.92 | 0.91 | 0.92 |

| | | | | | | |
|---|---|---|---|---|---|---|
| *BioKG2Vec_Concatenation_rf* | *0.91* | *0.94* | *0.92* | *0.91* | *0.91* | *0.92* |
| *DistMult_Average_xgb* | *0.91* | *0.94* | *0.92* | *0.91* | *0.90* | *0.92* |
| *DistMult_Sum_xgb* | *0.91* | *0.94* | *0.92* | *0.91* | *0.90* | *0.92* |
| *DistMult_Concatenation_svm* | *0.91* | *0.93* | *0.94* | *0.91* | *0.90* | *0.91* |
| *N2V_Sum_svm* | *0.90* | *0.94* | *0.91* | *0.90* | *0.90* | *0.92* |
| *DistMult_Hadmard_xgb* | *0.91* | *0.94* | *0.91* | *0.91* | *0.90* | *0.92* |
| *DistMult_Hadmard_rf* | *0.91* | *0.94* | *0.92* | *0.90* | *0.90* | *0.91* |
| *N2V_Concatenation_xgb* | *0.91* | *0.93* | *0.92* | *0.91* | *0.90* | *0.91* |
| *TransE_Average_ffn* | *0.91* | *0.93* | *0.93* | *0.91* | *0.90* | *0.91* |
| *DistMult_Concatenation_rf* | *0.91* | *0.93* | *0.93* | *0.91* | *0.90* | *0.91* |
| *DistMult_Sum_svm* | *0.91* | *0.93* | *0.94* | *0.91* | *0.90* | *0.91* |
| *TransE_Sum_ffn* | *0.91* | *0.93* | *0.93* | *0.91* | *0.90* | *0.91* |
| *DistMult_Average_rf* | *0.91* | *0.93* | *0.93* | *0.91* | *0.90* | *0.91* |
| *TransE_Sum_svm* | *0.90* | *0.93* | *0.92* | *0.90* | *0.90* | *0.91* |
| *TransE_Hadmard_rf* | *0.90* | *0.94* | *0.91* | *0.90* | *0.90* | *0.91* |
| *DistMult_Sum_rf* | *0.91* | *0.93* | *0.93* | *0.91* | *0.89* | *0.91* |
| *TransE_Sum_xgb* | *0.90* | *0.93* | *0.92* | *0.90* | *0.89* | *0.91* |
| *TransE_Average_xgb* | *0.90* | *0.93* | *0.92* | *0.90* | *0.89* | *0.91* |
| *BioKG2Vec_Average_xgb* | *0.89* | *0.94* | *0.89* | *0.89* | *0.89* | *0.91* |
| *BioKG2Vec_Sum_xgb* | *0.89* | *0.94* | *0.89* | *0.89* | *0.89* | *0.91* |
| *BioKG2Vec_Hadmard_svm* | *0.88* | *0.96* | *0.85* | *0.88* | *0.89* | *0.91* |
| *BioKG2Vec_Hadmard_xgb* | *0.88* | *0.96* | *0.85* | *0.88* | *0.89* | *0.91* |
| *BioKG2Vec_Hadmard_ffn* | *0.88* | *0.96* | *0.84* | *0.87* | *0.89* | *0.91* |
| *N2V_Average_svm* | *0.89* | *0.93* | *0.90* | *0.89* | *0.89* | *0.90* |
| *N2V_Average_ffn* | *0.89* | *0.93* | *0.90* | *0.89* | *0.88* | *0.90* |
| *N2V_Sum_ffn* | *0.89* | *0.92* | *0.91* | *0.89* | *0.88* | *0.90* |
| *TransE_Hadmard_lr* | *0.88* | *0.93* | *0.89* | *0.88* | *0.88* | *0.90* |
| *TransE_Hadmard_svm* | *0.89* | *0.93* | *0.90* | *0.89* | *0.88* | *0.90* |
| *DistMult_Average_svm* | *0.90* | *0.91* | *0.94* | *0.90* | *0.88* | *0.89* |
| *N2V_Concatenation_rf* | *0.89* | *0.91* | *0.93* | *0.89* | *0.88* | *0.89* |
| *TransE_Average_rf* | *0.90* | *0.91* | *0.94* | *0.90* | *0.88* | *0.89* |
| *TransE_Sum_rf* | *0.90* | *0.91* | *0.94* | *0.90* | *0.88* | *0.89* |
| *DistMult_Hadmard_lr* | *0.89* | *0.92* | *0.91* | *0.89* | *0.88* | *0.89* |
| *TransE_Average_svm* | *0.89* | *0.91* | *0.91* | *0.89* | *0.87* | *0.89* |
| *DistMult_Hadmard_svm* | *0.89* | *0.91* | *0.91* | *0.88* | *0.87* | *0.89* |
| *BioKG2Vec_Hadmard_rf* | *0.87* | *0.93* | *0.86* | *0.87* | *0.87* | *0.90* |
| *DistMult_Concatenation_lr* | *0.88* | *0.91* | *0.92* | *0.89* | *0.87* | *0.89* |
| *N2V_Sum_xgb* | *0.88* | *0.92* | *0.90* | *0.88* | *0.87* | *0.89* |
| *N2V_Average_xgb* | *0.88* | *0.92* | *0.90* | *0.88* | *0.87* | *0.89* |
| *BioKG2Vec_Concatenation_lr* | *0.88* | *0.91* | *0.91* | *0.88* | *0.87* | *0.89* |
| *TransE_Concatenation_lr* | *0.88* | *0.90* | *0.93* | *0.89* | *0.87* | *0.88* |
| *DLEMB_Concatenation_lr* | *0.88* | *0.90* | *0.92* | *0.88* | *0.86* | *0.88* |

| | | | | | | |
|---|---|---|---|---|---|---|
| *N2V_Hadmard_ffn* | *0.87* | *0.92* | *0.88* | *0.87* | *0.86* | *0.89* |
| *DistMult_Sum_lr* | *0.88* | *0.90* | *0.91* | *0.88* | *0.86* | *0.88* |
| *DistMult_Average_lr* | *0.88* | *0.90* | *0.92* | *0.88* | *0.86* | *0.88* |
| *N2V_Hadmard_xgb* | *0.86* | *0.92* | *0.87* | *0.86* | *0.86* | *0.88* |
| *N2V_Hadmard_svm* | *0.85* | *0.92* | *0.85* | *0.85* | *0.86* | *0.88* |
| *BioKG2Vec_Average_lr* | *0.87* | *0.90* | *0.90* | *0.87* | *0.85* | *0.87* |
| *BioKG2Vec_Sum_lr* | *0.87* | *0.90* | *0.90* | *0.87* | *0.85* | *0.87* |
| *BioKG2Vec_SAGE_Concatenation_xgb* | *0.87* | *0.89* | *0.91* | *0.87* | *0.85* | *0.87* |
| *BioKG2Vec_Hadmard_lr* | *0.84* | *0.93* | *0.81* | *0.84* | *0.85* | *0.88* |
| *N2V_Concatenation_lr* | *0.86* | *0.89* | *0.90* | *0.86* | *0.84* | *0.87* |
| *N2V_Hadmard_lr* | *0.84* | *0.92* | *0.83* | *0.84* | *0.84* | *0.87* |
| *BioKG2Vec_Sum_rf* | *0.86* | *0.88* | *0.91* | *0.86* | *0.84* | *0.86* |
| *BioKG2Vec_SAGE_Concatenation_rf* | *0.86* | *0.89* | *0.90* | *0.86* | *0.84* | *0.86* |
| *BioKG2Vec_Average_rf* | *0.86* | *0.88* | *0.91* | *0.86* | *0.84* | *0.86* |
| *N2V_Average_rf* | *0.86* | *0.88* | *0.91* | *0.86* | *0.83* | *0.86* |
| *BioKG2Vec_GCN_Concatenation_xgb* | *0.86* | *0.87* | *0.91* | *0.86* | *0.83* | *0.86* |
| *N2V_Sum_rf* | *0.86* | *0.87* | *0.91* | *0.86* | *0.83* | *0.86* |
| *DLEMB_Sum_lr* | *0.85* | *0.87* | *0.90* | *0.85* | *0.83* | *0.85* |
| *DLEMB_Average_lr* | *0.85* | *0.87* | *0.90* | *0.85* | *0.83* | *0.85* |
| *N2V_Hadmard_rf* | *0.84* | *0.88* | *0.89* | *0.84* | *0.82* | *0.85* |
| *BioKG2Vec_GCN_Concatenation_rf* | *0.84* | *0.88* | *0.88* | *0.84* | *0.82* | *0.85* |
| *N2V_Sum_lr* | *0.84* | *0.88* | *0.89* | *0.84* | *0.82* | *0.85* |
| *N2V_Average_lr* | *0.84* | *0.88* | *0.89* | *0.84* | *0.82* | *0.85* |
| *TransE_Average_lr* | *0.82* | *0.85* | *0.89* | *0.83* | *0.80* | *0.83* |
| *TransE_Sum_lr* | *0.82* | *0.85* | *0.89* | *0.83* | *0.80* | *0.83* |
| *BioKG2Vec_GCN_Concatenation_ffn* | *0.80* | *0.83* | *0.87* | *0.80* | *0.77* | *0.81* |
| *BioKG2Vec_GCN_Sum_rf* | *0.79* | *0.83* | *0.86* | *0.79* | *0.76* | *0.80* |
| *BioKG2Vec_GCN_Average_rf* | *0.79* | *0.83* | *0.86* | *0.79* | *0.76* | *0.80* |
| *BioKG2Vec_GCN_Sum_ffn* | *0.77* | *0.81* | *0.87* | *0.78* | *0.74* | *0.79* |
| *BioKG2Vec_GCN_Average_ffn* | *0.77* | *0.80* | *0.89* | *0.78* | *0.73* | *0.78* |
| *BioKG2Vec_GCN_Sum_xgb* | *0.76* | *0.80* | *0.87* | *0.77* | *0.72* | *0.78* |
| *BioKG2Vec_GCN_Average_xgb* | *0.76* | *0.80* | *0.87* | *0.77* | *0.72* | *0.78* |
| *BioKG2Vec_GCN_Hadmard_rf* | *0.76* | *0.80* | *0.87* | *0.77* | *0.72* | *0.78* |
| *BioKG2Vec_GCN_Concatenation_svm* | *0.77* | *0.79* | *0.90* | *0.78* | *0.72* | *0.77* |
| *BioKG2Vec_GCN_Hadmard_xgb* | *0.75* | *0.78* | *0.87* | *0.76* | *0.70* | *0.77* |
| *BioKG2Vec_GCN_Sum_svm* | *0.75* | *0.78* | *0.89* | *0.76* | *0.70* | *0.77* |
| *BioKG2Vec_GCN_Average_svm* | *0.73* | *0.77* | *0.88* | *0.75* | *0.68* | *0.75* |
| *BioKG2Vec_GCN_Hadmard_ffn* | *0.74* | *0.76* | *0.91* | *0.75* | *0.68* | *0.75* |
| *BioKG2Vec_GCN_Average_lr* | *0.71* | *0.75* | *0.90* | *0.73* | *0.66* | *0.74* |
| *BioKG2Vec_GCN_Sum_lr* | *0.71* | *0.75* | *0.90* | *0.73* | *0.66* | *0.74* |
| *BioKG2Vec_GCN_Concatenation_lr* | *0.71* | *0.75* | *0.89* | *0.73* | *0.65* | *0.74* |
| *BioKG2Vec_GCN_Hadmard_lr* | *0.61* | *0.68* | *0.96* | *0.68* | *0.56* | *0.68* |

| | | | | | | |
|---|---|---|---|---|---|---|
| *BioKG2Vec_GCN_Hadmard_svm* | *0.52* | *0.66* | *1.00* | *0.66* | *0.50* | *0.66* |

*Supplementary table 2: Results of 5-fold cross-validation on randomly selected diseases belonging to different number of GDA stratifications (algorithm is support vector machine, and GDA representation is concatenation). N_ass represents ….*

| | RANDOM | | | | | | DLEMB | | | | |
|---|---|---|---|---|---|---|---|---|---|---|---|
| **CUI** | **n_ass** | **mean_accuracy** | **mean_precision** | **mean_recall** | **mean_f1** | **mean_roc** | **mean_accuracy** | **mean_precision** | **mean_recall** | **mean_f1** | **mean_roc** |
| **C0003865** | 40 | 0.41 | 0.42 | 0.45 | 0.42 | 0.43 | 0.9 | 0.91 | 0.9 | 0.9 | 0.97 |
| **C0920563** | 60 | 0.51 | 0.5 | 0.52 | 0.51 | 0.51 | 0.89 | 0.86 | 0.97 | 0.9 | 0.97 |
| **C0004096** | 80 | 0.42 | 0.41 | 0.41 | 0.41 | 0.46 | 0.81 | 0.8 | 0.85 | 0.82 | 0.88 |
| **C0234533** | 101 | 0.46 | 0.46 | 0.46 | 0.45 | 0.48 | 0.95 | 0.95 | 0.95 | 0.95 | 0.99 |
| **C0236736** | 117 | 0.52 | 0.52 | 0.5 | 0.51 | 0.53 | 0.89 | 0.88 | 0.89 | 0.89 | 0.95 |
| **C0005684** | 141 | 0.5 | 0.5 | 0.49 | 0.49 | 0.5 | 0.86 | 0.85 | 0.87 | 0.86 | 0.92 |
| **C0269102** | 161 | 0.46 | 0.46 | 0.45 | 0.45 | 0.46 | 0.76 | 0.76 | 0.76 | 0.76 | 0.85 |
| **C0151744** | 176 | 0.48 | 0.48 | 0.49 | 0.48 | 0.48 | 0.77 | 0.75 | 0.81 | 0.78 | 0.84 |
| **C0020538** | 190 | 0.49 | 0.49 | 0.51 | 0.5 | 0.51 | 0.86 | 0.85 | 0.87 | 0.86 | 0.93 |
| **C0011860** | 221 | 0.47 | 0.47 | 0.45 | 0.45 | 0.48 | 0.82 | 0.82 | 0.83 | 0.82 | 0.88 |
| **C0004352** | 261 | 0.5 | 0.5 | 0.48 | 0.48 | 0.52 | 0.78 | 0.79 | 0.77 | 0.77 | 0.87 |
| **C0009404** | 277 | 0.5 | 0.49 | 0.5 | 0.49 | 0.52 | 0.79 | 0.78 | 0.81 | 0.79 | 0.86 |
| **C0024623** | 300 | 0.5 | 0.5 | 0.5 | 0.5 | 0.5 | 0.78 | 0.76 | 0.81 | 0.79 | 0.86 |
| **C3658290** | 404 | 0.44 | 0.44 | 0.44 | 0.44 | 0.46 | 0.92 | 0.9 | 0.94 | 0.92 | 0.96 |
| **C3714756** | 447 | 0.45 | 0.45 | 0.44 | 0.44 | 0.43 | 0.87 | 0.84 | 0.91 | 0.87 | 0.91 |
| **C0005586** | 477 | 0.5 | 0.5 | 0.53 | 0.52 | 0.5 | 0.79 | 0.78 | 0.81 | 0.8 | 0.87 |
| **C1257931** | 525 | 0.51 | 0.51 | 0.49 | 0.5 | 0.51 | 0.83 | 0.8 | 0.89 | 0.84 | 0.9 |
| **C0678222** | 538 | 0.52 | 0.52 | 0.52 | 0.52 | 0.53 | 0.86 | 0.83 | 0.91 | 0.87 | 0.93 |
| **C0033578** | 616 | 0.51 | 0.51 | 0.52 | 0.51 | 0.51 | 0.77 | 0.76 | 0.8 | 0.78 | 0.84 |
| **C0009402** | 702 | 0.49 | 0.49 | 0.48 | 0.49 | 0.48 | 0.69 | 0.68 | 0.72 | 0.7 | 0.75 |
| **C0023893** | 774 | 0.51 | 0.51 | 0.52 | 0.51 | 0.49 | 0.71 | 0.7 | 0.74 | 0.72 | 0.77 |
| **C0036341** | 883 | 0.51 | 0.51 | 0.53 | 0.52 | 0.51 | 0.75 | 0.75 | 0.76 | 0.75 | 0.83 |
| **C0006142** | 1074 | 0.48 | 0.48 | 0.49 | 0.49 | 0.46 | 0.72 | 0.7 | 0.76 | 0.73 | 0.78 |
